# Applying non-parametric testing to discrete transfer entropy

**DOI:** 10.1101/460733

**Authors:** Wilkie Olin-Ammentorp, Nathaniel Cady

## Abstract

Transfer entropy (TE) is a powerful algorithm which attempts to detect the transfer of information from one system to another. In neuroscience, it has the potential to track the movement of information through complex neuronal systems, and provide powerful insights into their organization and operation. One such application is the ability to infer the existence of causal connectivity (such as synaptic pathways) between neurons in a culture being recorded by micro-electrode array (MEA).

There are several challenges, however, in applying TE to neurological data; one of these is the ability to robustly classify what experimental TE value qualifies as significant. We find that common methods in spike train analysis such as a Z-test cannot be applied, as their assumptions are not met. Instead, we utilize surrogate data to compute a sample under the null hypothesis (no causal connection), and resample experimental data through Markov chain Monte Carlo (MCMC) methods to create a sample of TE values under experimental conditions. A standard non-parametric test (Mann-Whitney U-test) is then applied to compare these samples, and determine if they represent a significant connection.

We have applied this methodology to MEA recordings of neuronal cultures developing over a period of roughly a month, and find that it provides a wealth of information regarding the cultures' maturity. This includes features such as the directed graph of causal connections across the MEA and identification of information exchange centers. These results are consistent and carry a well-defined significance level.

## Introduction

Micro-electrode arrays (MEAs) are tools which contain microscopic electrodes, allowing for measurements of electrical activity across a variety of samples (such as cultures of primary neurons). The resolution and capabilities of MEAs are growing, as measurement electrodes become smaller, more numerous, and more sensitive ([1]). While this allows researchers to study the development and behavior of neuronal networks in unprecedented detail, it also creates challenges in data analysis as the size and complexity of experimental results grows. To develop a clearer understanding of the connectivity of neuronal tissue, algorithms for data analysis must provide results with a high confidence, and have common, efficient implementations available freely.

The focus of many MEA experiments is to determine the functional connectivity between the neurons, and to investigate how this connectivity develops or changes under differing conditions ([2-6]). Many different mathematical methods have been applied to infer functional connectivity from the spiking activity of neurons, including cross-correlation (CC) and transfer entropy (TE) ([7, 8]).

A challenge for both these algorithms is determining what value is significant, as both methods have biases and can be fooled by coincidences in the dataset, producing non-zero values even when a significant connection does not exist ([9], [10]). As a result, it is necessary to filter these data using estimates of their values under the null hypothesis that no connection exists.

We present a novel Markov-chain Monte Carlo (MCMC) based method of detecting significance from TE results, by sampling surrogate and experimental values. A non-parametric statistical test, the Mann-Whitney U-test, is then applied to these samples to determine the significance of each hypothesis regarding the connectivity.

We then apply this method to analyze the connectivity found within experimental recordings of developing neuronal cultures created by Wagenaar et al. ([11]). The TE results provide a wealth of information, including the undirected graph of connectivity during development, and the relative importance of different sites in exchanging information.

Lastly, we briefly the performance of our software, written in the Julia language and compatible with common Python tools for plotting, network analysis, and browser-based interactivity [12-14]. As a result, our implementation is fully open-source, portable, and can be quickly built from an image.

## 1 Methods

### 1.1 Transfer Entropy

Transfer entropy (TE) is an information-theory based measure which quantifies how much one’s prediction of a signal changes based on the state of another signal ([15]). It is a measure which has been widely adopted in neuroscience ([10, 16]), as it has been found to have give the most accurate results from several different methods applied to simulated neural networks [17].

As detailed by Vicente et al. [10], TE may be calculated on discrete data (such as spike trains) by creating a ‘delay vector.’ Given an original, discrete spike train *x*(*t*) which can indicate the presence or absence of a spike at *t*, let ***x***(*w*) be a vector counting the number of spikes in each temporal bin of width *w*, with sufficient bins to span *x*(*t*). The short-term behavior of ***x***(*w*) at an index *i* is then represented a single state, by creating a delay vector 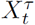, which captures ***x***_*i*_ from *i* to the previous *τ* — 1 values.

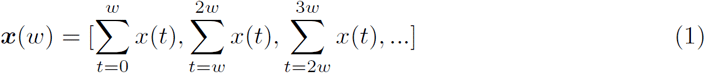

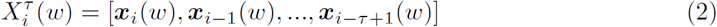

The combination of a signal *x*’s past state, its next value, and the past state of another signal *y* are used to create a joint distribution (JD, Fig 1a). The objective of this JD is to summarize the probability of each future state of *x*, given the delay vectors representing the past of both signals. By summing this JD along its rows, a marginal distribution (MD) is found, which predicts the future of *x* using only its own past state, excluding the past of *y*. Often, the past state may be offset by an additional *o* bins, inspecting for relationships which may have a delay.

The divergence between conditional probabilities predicting the future of *x* using the JD and MD is the transfer entropy between the signals. When the information from *y* is useful to predict the future of *x*, the divergence between these distributions will increase, indicating the possibility of a causal relationship from *y* to *x*. The inverse is not true, as TE is not commutative; the influence of *x* on *y* must be separately examined.

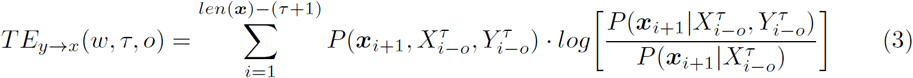

While TE is a powerful informational method for investigating possible causality between different signals, it is computationally expensive. To apply it to spiking data, one must compute joint distributions of all spike embeddings between all combinations of electrodes on the array. One must also select the dimension of the delay embedding (*τ*), and the period the delay embedding spans. The correct values for these parameters are not obvious, and each has a significant impact on the algorithm’s end result. Additionally, it has been shown that TE only achieves peak accuracy for spike trains when it searches for causality at multiple time-offsets, as neurons influence one another at differing time scales ([18]).

**Fig.1.**
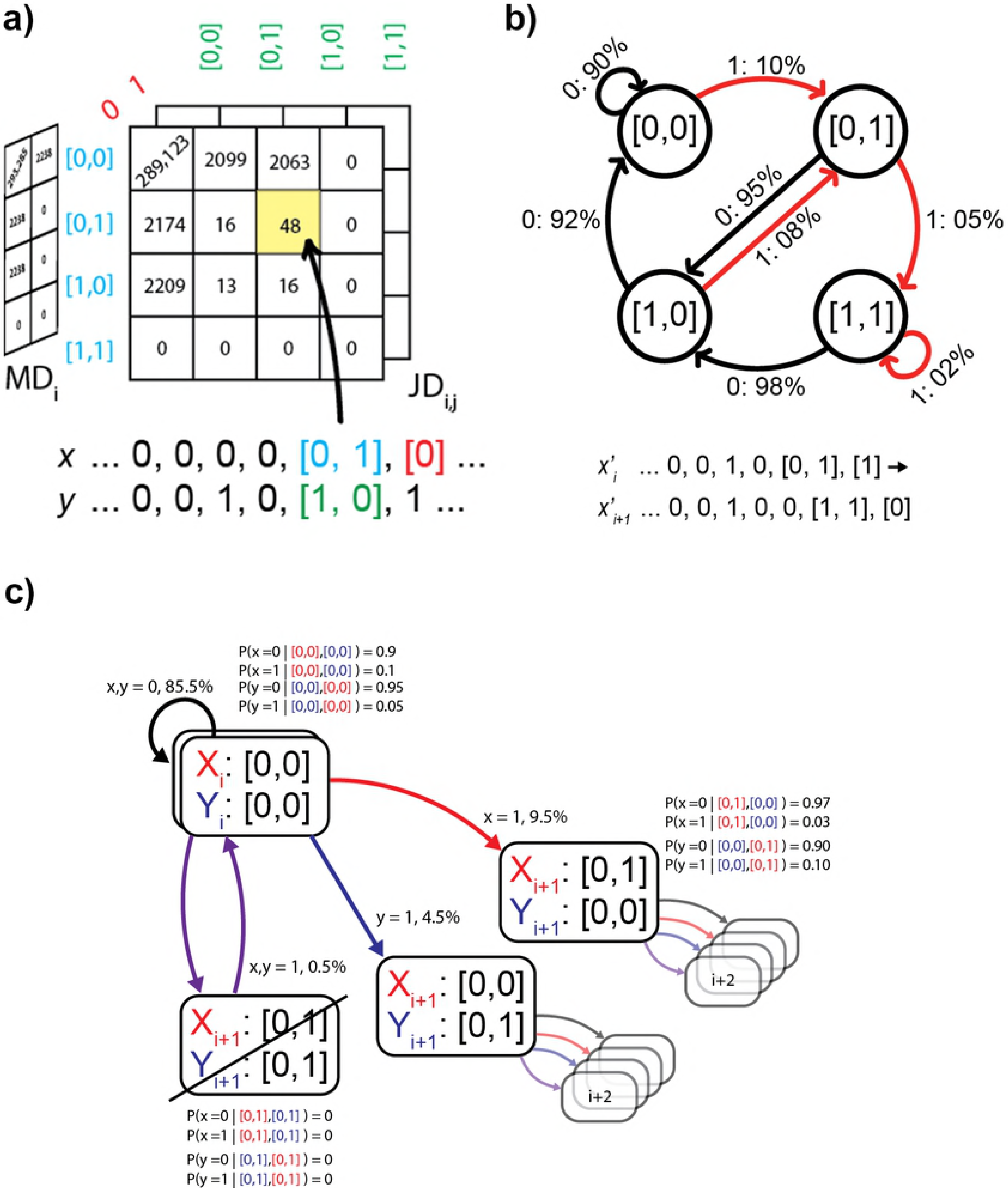
Illustration of calculating TE. a) Illustration of generating the joint distribution (JD) which describes the future state of a signal given its past state, and the past state of another signal. It can be reduced to the marginal distribution (MD, left side), which predicts its future outcome based on only its own past state. The divergence between each distribution's predictions is used to calculate TE. b) Markov-chain Monte Carlo (MCMC) method of generating spike count vectors representing the null hypothesis (no connectivity). In this example, the embedding length *τ* is 2, and the maximum possible spike count in a bin is 1. This creates a simple system which can transition through 4 states to generate a spike train. c) The MCMC method of resampling experimental TE values. Two signals' futures must be chosen randomly using the conditional probabilities from their original JDs. This provides a transition to the next joint state. If this state was not encountered in the original recording, it must be rejected (bottom).

### 1.2 Significance Testing

Both TE and other algorithms generally analyze entire MEA recordings as a single sample. As a result, only one experimental value is calculated for each possible relation between spiking sources. Due to coincidental behaviors, noise, and biases which may be present in the recordings and algorithms, TE values are almost always non-zero.

To determine what level a potential relation must exceed in order to be significant, it must be compared to the amount which could be expected under the null hypothesis (there is no causal relationship between the sources). If a value exceeds this, the alternative hypothesis (there exists a causal connection between the sources) is accepted.

#### 1.2.1 Traditional Approaches

‘Surrogate’ datasets approximating each spike train operating under the null hypothesis of no connectivity can be created by several methods. All of these methods must tread a thin path between maintaining the overall properties of a recording (such as spike count and inter-spike intervals), while obfuscating information that was created by causal relationships between sources ([9]).

Most often, this is done by shuffling data, exchanging spikes, or dithering times in the original recording. This is done many times to create a sample of results under the null hypothesis that there is no connectivity. For cross-correlation and other methods, these values can be well-described by a normal distribution, allowing parametric methods such as the Z-test to be applied. If the original experimental value exceeds a certain sigma level (typically 3-6*σ*) of the distribution, it can be accepted as significant with the well-defined risks corresponding to the test being used ([19]).

However, the complex and non-linear behavior of TE complicates statistical testing of its values' significance. This can lead to assumptions of standard parametric tests being violated, making appropriate application of these tests challenging or impossible.

Non-parametric techniques offer an alternative approach to calculating the significance, while making fewer assumptions about the underlying distributions. If a large enough sample of values can be found for each hypothesis, a variety of non-parametric methods can be applied.

#### 1.2.2 MCMC Approach for TE

The unique way in which TE captures the dynamics of signal pairs allows it to be resampled under both the null and alternative hypotheses. Specifically, the joint and marginal distributions calculated from a pair of signals in the recording can be used to create Markov chains, which can be used in a manner similar to Gibbs sampling, to calculate values under either hypothesis ([20]).

Under the simpler case of the null hypothesis, only one MD for each source is needed. Assuming a source starts at a quiet state, its first state vector 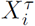 is created. Using the conditional probability 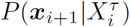, an outcome for the next state ***x***_*i*+1_ is randomly selected. The state vector is then moved forward in time to include this outcome, and this process is repeated until a sample of sufficient length is created (Fig 1b). In our implementation, spike-count vectors for each channel are generated until the total number of spikes is equal to that of the original recording. This surrogate recording is then re-analyzed to calculate TE values for each possible connection. In this case, each source is independently sampled from a conditional distribution dependent on only its own past, and there is no causal behavior between sources. As a result, any detected information transfer is a due to the TE algorithm's biases.

The more challenging case is to re-sample the value of each possible connection under the alternative hypothesis that sources do influence one another. In this case, the joint distributions predicting the next state of the two relevant sources must be simultaneously sampled. Similarly to the null case, this is done by starting at a quiet state, and randomly choosing each signal's next state based on the conditional probability 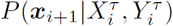, given the joint state between both sources’ state vectors. The occurrence of each new joint state is incremented in a JD, and the process is repeated until the number of total joint states in this distribution equals the number calculated using the original recording.

One complication arises, however: both signals are being generated at once, and they can occasionally choose conflicting states for the next step. This represents a state which was never encountered in the original recording (such as both channels spiking simultaneously). When this rare situation occurs, outcomes are re-chosen until a new, valid configuration is found. At least one such configuration always exists; given that the signal entered one state in phase space, it also had to traverse out of it (the trajectory of the signal is closed, given that the beginning and ending of each recording corresponds to a quiescent state).

Additionally, these Markov chains can only be made from distributions which were created at a zero offset. When the next outcome is delayed from the current state vector, the next state vector is not immediately dependent on the previous spiking state, and a Markov chain which gives appropriate state-changes cannot be constructed. As a result, to re-sample TE values at a non-zero delay, the signal must be reconstructed from the 0-delay distribution, and the delayed state measured from its generated signal.

Once each possible connection has had its JD resampled, a new TE value for this connection can be calculated. Essentially, the difference between this re-sampled TE value and the original experimental value represents the result’s sensitivity to slight perturbations to, or recombinations of the original signals. If the original value depended solely on a handful of extraordinary events, the spread of resampled signals will be large, indicating uncertainty.

The spread of this value is important when estimating the significance of the alternative hypothesis compared to the null. A single experimental value may always exceed the surrogate values under the null hypothesis, but re-sampling these experimental values reveals that there may be significant overlap between the samples.

To test the null hypothesis, a direct Mann-Whitney U-test is applied. The U-test is a standard, unpaired, non-parametric test which compares all possible matches between the surrogate and experimental samples. It examines the proportion of matches in which the experimental samples are lesser. If this proportion is below the risk level, the null hypothesis is rejected, and the difference between these two samples indicates that the experimental TE value is significant, and provides sufficient evidence to infer a causal connection exists.

We denote this method as ‘resampling transfer entropy’ (R-TE), as it compares surrogate data and resampled experimental values to determine if a causal connection is significant.

### 1.3 Implementation

Julia ([12]) is a recent programming language with a focus on usability, parallelism, and numerical performance. This gives it number of advantages: interactivity, integration with Python libraries, and parallel dispatch. Additionally, Julia is free and open-source software, aiming to provide a common base for scientific computing. Because of these strengths, we use it to implement R-TE and additional functions needed for data handling, analysis, and statistical testing. As it is completely written on a free and open-source base, our software can be easily downloaded, built, and executed inside a container-based system ([21]).

The Jupyter notebook system ([14]) supports Julia kernels, allowing analysis commands to be carried out interactively through a browser. Results can then be explored and displayed in-line, by utilities such as Matplotlib ([22]), and this methodology was used to generate most the figures which we present. One exception is representations of graphs (Fig 5d-f), which were generated using the Gephi software ([23]).

The Julia language is supported on all common operating systems (Windows, Macintosh, Linux), and we execute our software on an Ubuntu 14.04-based system with 2 Xeon E5-2670 CPUs (2.6 GHz, 32 total threads). Runtime data was collected using macros included in the Julia language.

### 1.4 Experimental Data Source

To test the viability and performance of R-TE on experimental data, we applied it to MEA recordings created by Wagenaar et al., as they provide a large collection of neuronal cultures which developed over several weeks [11]. These cultures were prepared under differing conditions, including the source of the cortical material and the density at which it was plated. Cortical material was collected from eight Wistar rat embryos, giving rise to 8 ‘batches’ of source material. The material was then disassociated, and plated into individual cultures on MEAs. After 3-4 days *in vitro* (DIV), daily recordings were taken of the spiking activity as measured by the 59 recording electrodes on the array.

We sub-selected 4 cultures to investigate as a demonstration of R-TE’s capabilities. The first two cultures, A and B, were densely plated (2.5 ± 1.5*x*10^3^ cells/mm^2^ - batch 1, dense cultures 1 and 2). Culture C has a ‘small’ plating density, with 1.6 ± 0.6*x*10^3^cells/mm^2^ (batch 6, small culture 1), and culture D has a ‘sparse’ 0.6 ± 0.24*x*10^3^cells/mm^2^ plating density (batch 6, sparse culture 1). It was reported that the sparser cultures C and D developed more slowly than the dense cultures A and B, as the former required more time to develop complex spiking features. This contrast provides an interesting case to apply R-TE, and compare the developmental kinetics between conditions.

## 2 Results & Discussion

### 2.1 Signal Dynamics

Obtaining meaningful results from TE is heavily dependent on selecting parameters which allow it to accurately capture the dynamical behavior of the system being investigated. For continuous signals, standard tests such as the Cao or Ragwitz criteria can be applied to find the proper parameters ([24]). However, these criteria do not extend well to discrete systems, such as spike trains.

To find the appropriate bin width *w*, delay embedding length *τ*, and range of delay offsets *o*, we first make a basic inspection of the signal dynamics. A correlogram shows that many of the interactions between signals fall within a window of 20 ms (Fig 2a), providing a relevant period which the series of offsets *o* should inspect. Next, the minimum value of *w* is given by the sampling rate of the recording (25 kHz). At this minimum of *w* = 40 μs, no more than one spike can occur in each bin. As *w* increases, more than one spike can fall in each bin. By changing *w*, the maximum number of spikes within one bin increases. If the bin is too large, many spikes which could be interacting or transferring information will fall into a single bin, improperly capturing the signal. We see an inflection above 1 ms, where the maximum number of spikes begins to rapidly increase - this suggests that *w* should not be larger than this value (Fig 2b). With *o* and *w* selected from signal dynamics, *τ* must be selected from its effect on results. At larger values of *τ* and *w*, the improper embedding of the signal causes the JD to effectively begin ‘memorizing’ it, creating artificially high TE values. We find that *τ* = 3 is a good selection, as the results with a larger or smaller *τ* do not differ strongly (Fig 2c). When results are analyzed, the power of the test being applied also depends on the number of resampled and surrogate values being sampled. Sampling the variation in total number of significant connections detected by bootstrap, we find that 32 samples of each is more than sufficient to effectively eliminate the variation in a tests’s outcome (Fig 2d). These parameters (*w* = 1 ms, *τ* = 3, *o* = 0-20 bins, 32 resamples/surrogates) are used to analyze the recordings for cultures A-D.

### 2.2 Comparison to Standard Methods

Each culture originates from a sample of disassociated neurons, which have no connections when they are first plated onto the MEA. This gives a reference starting point for algorithms detecting connectivity; early in a culture’s development, the number of detected causal connections should be either very small or zero.

This expected result, however, is not found when attempting to use a Z-test to estimate significance of experimental TE values. Even when using what would be an extremely high significance level if the test’s assumptions were met, the test fails to reject hundreds or thousands of connections which must be insignificant, given the known developmental status of the culture. In contrast, R-TE quickly begins to filter out the majority of these insignificant connections when standard significance levels (*a* ≤5%) are used (Fig 3a). But as the cultures develop, R-TE begins to detect connections with a significance which cannot be reasonably rejected (Fig 3b).

**Fig.2.**
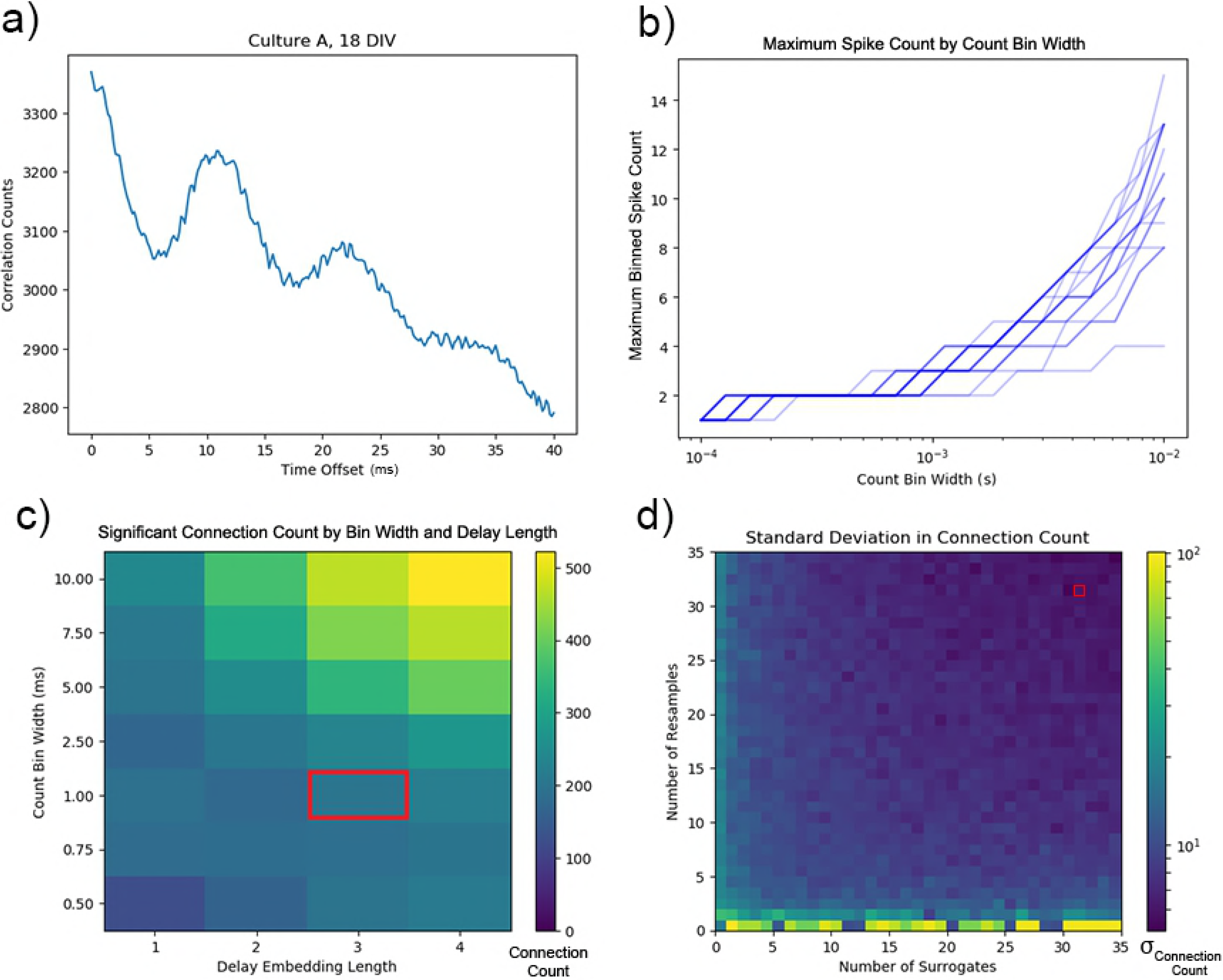
Parameter Selection for TE. a) Total correlation counts between pairs of signals over a range of delays (Culture A, 25 days *in vitro* (DIV)). Many of the strongest interactions occur within 20 ms. b) The maximum number of spike counts found in a single bin, for recordings from 8-18 DIV for culture A. After a bin width of 1 ms, many events can begin falling within a single bin, indicating the signal is being improperly embedded. c) The total number of significant connections inferred by R-TE while varying *w* and *τ*, given a constant *o* which ranges from 0-20 bins offset (Culture C, 28 DIV). A region of parameters which leads to results that do not change greatly indicates possible good values to use. The selected parameters *w* =1 ms, *τ*= 3 are outlined in red. d) The standard deviation in number of significant connections for a sample file (culture A, 10 DIV), by the number of resampled and surrogate values. The selected parameter (32 of each) is highlighted in red.

**Fig.3.**
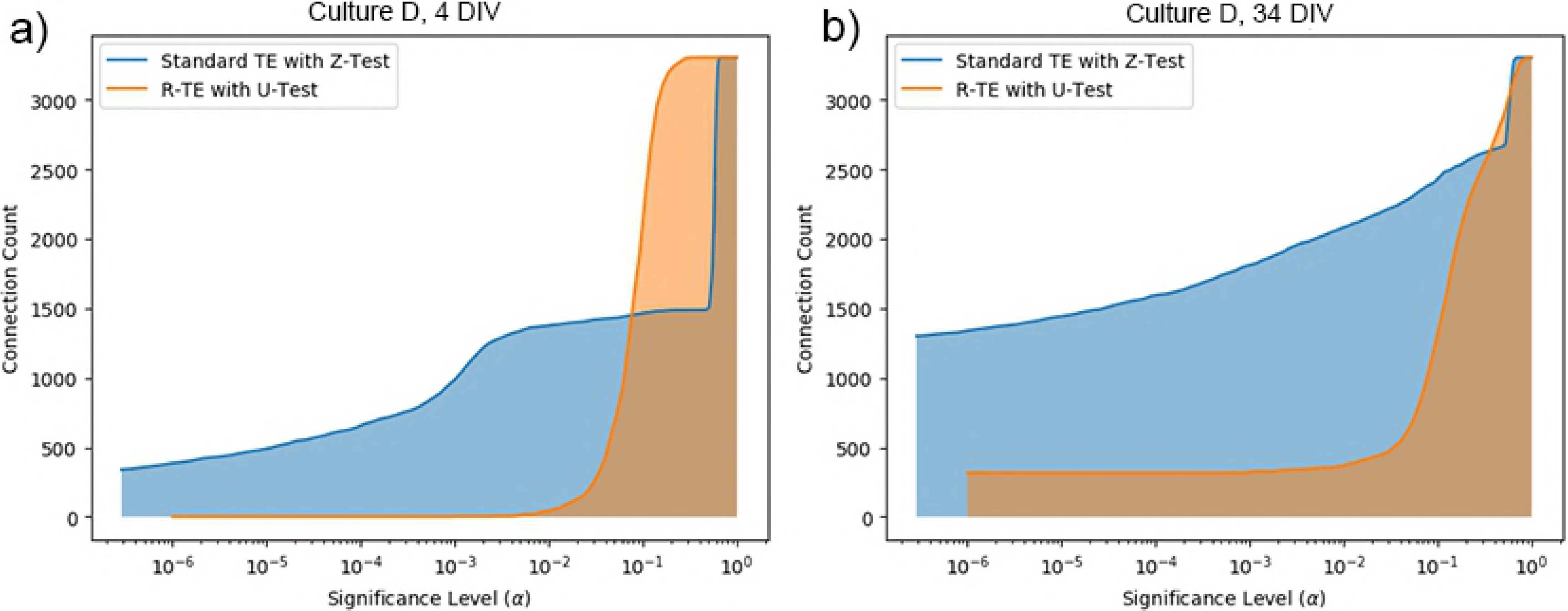
Comparing the Z-test and U-test for TE. a) The number of significant connections inferred by standard TE versus R-TE by significance level, early in the development (4 DIV) of the sparse culture, D. At each level, the number of rejected connections are above each line, and accepted are below. b) The same plot for a culture D at a later stage of maturity (34 DIV).

Samples of TE values from surrogate and resampled experimental data show that the Z-test's lack of discriminative power here is a result of samples being poorly described by Gaussians. This is particularly the case in early days where their values are low (Fig 4a,b). As development progresses and values grow, Gaussians may become suitable (Fig 4c), but it is not a safe assumption to make for all cases. This motivates our non-parametric approach to testing significance.

**Fig.4.**
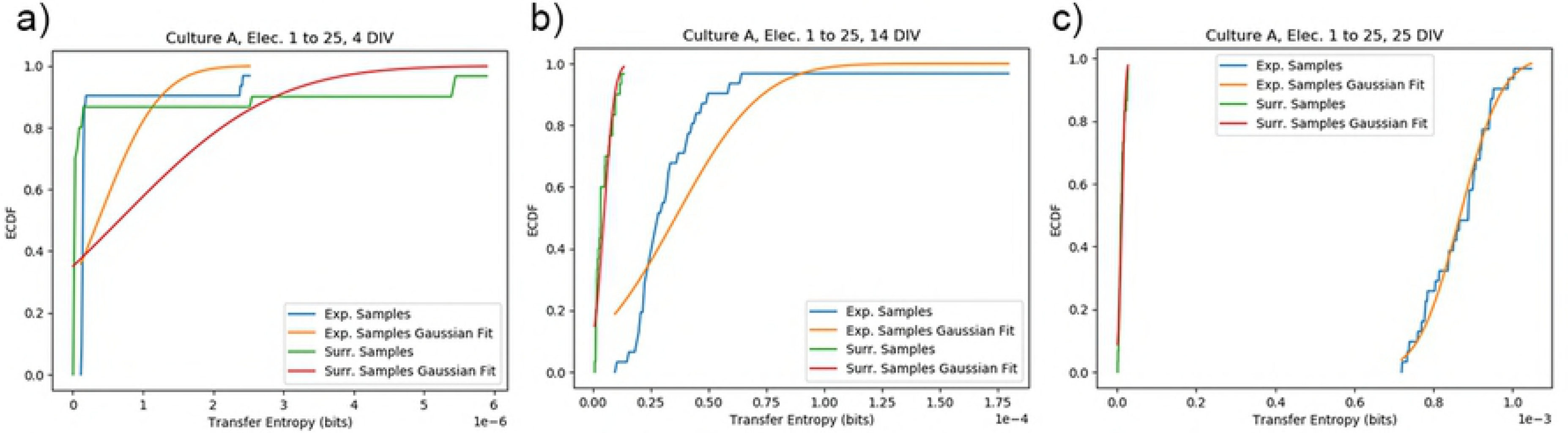
The normality of TE values. a) Samples of TE values from surrogate and re-sampled experimental data are shown with Gaussian distributions which attempt to describe them. Slight perturbations to JDs at this stage can cause relatively large shifts in TE values. b) The same plot after an additional 10 DIV. TE values have grown, but are still not very well-described by a Gaussian. This does not occur until 25 DIV (c), when the large number of joint events causes TE values to only slightly change under perturbation.

### 2.3 Graphical Analysis of Development

All recordings of cultures A-D were analyzed using R-TE, with a significance threshold of *α*=1%. This yields an adjacency matrix for each recording, representing the connectivity of the neural network between the recording sites on the MEA (Fig 5a). As expected, the two dense cultures A and B show similar development, reaching near full connectivity after approximately two weeks *in vitro*. In contrast, cultures C and D (with small and sparse densities, respectively) develop more slowly and have fewer connections, as would be expected from their lower plating densities (Fig 5b). Additionally, the number of detected connections for all cultures does not vary greatly when standard significance levels (*α* ≤ 5%) are used (Fig 5c).

This finding is consistent with those of Wagenaar et al., who reported that cultures with lower plating densities developed more slowly and exhibited bursting behavior later than dense cultures. One feature of note is that the onset of greater connectivity levels in cultures A-D (Fig 5b) correlates to the onset of more consistent and complex bursting activity, as described in their original work ([11]).

**Fig.5.**
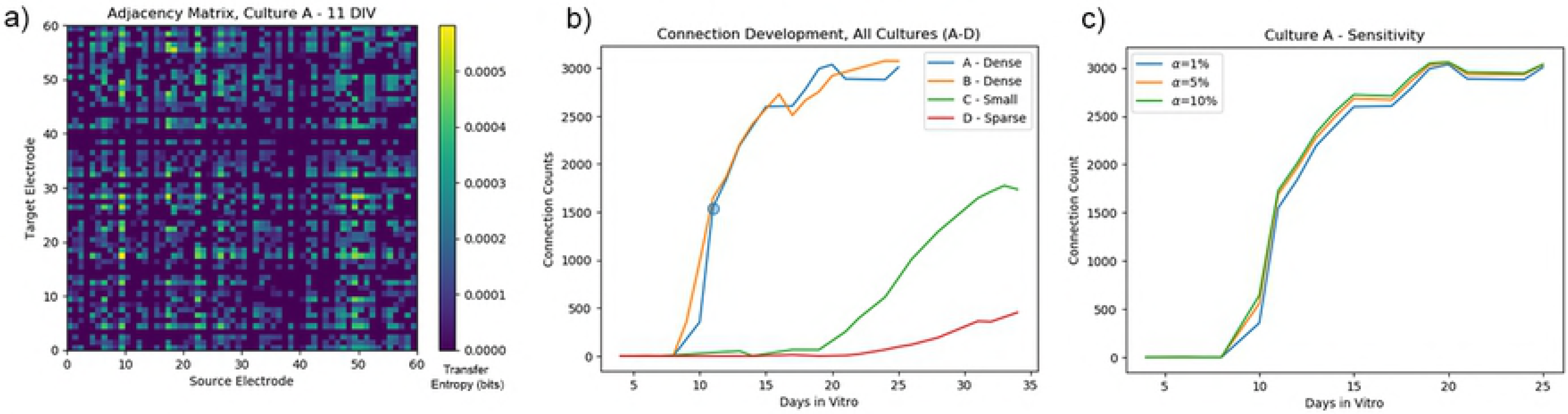
The development of neuronal connections, as detected by R-TE. a) The resulting adjacency matrix (with corresponding TE values) for culture A after 10 DIV. b) The total number of significant causal connections detected for each culture, by DIV. Denser cultures develop more connections more quickly than the small and sparse cultures. The blue point corresponds to the sum of all connectivity shown in subfigure a. c) The variation in number of accepted connections for this culture by significance level.

The TE values of each connection also provide insight into the structure of the neuronal network. Each connection's TE value approximates the exchange of information from one site to the other, adding a meaningful weight to each edge between nodes in the graph describing connectivity. By summing the TE values on edges incident on a node, the relative importance of this node in processing information can be seen.

We find that stable rankings of nodes, sorted by the sum of incoming TE, emerge alongside connectivity for all cultures (Fig 6a). In other words, connections which develop early remain important, and often grow in strength as the neuronal network's connectivity increases. (6d,e,f) Incoming TE also correlates well with a coarse description of centrality, the in-degree (Fig 5e,f). However, it provides more granular detail and does not saturate to the total number of nodes if the graph is fully connected. This suggests that incoming TE provides a metric for determining the relative importance of nodes, and identifying centers of information processing.

**Fig.6.**
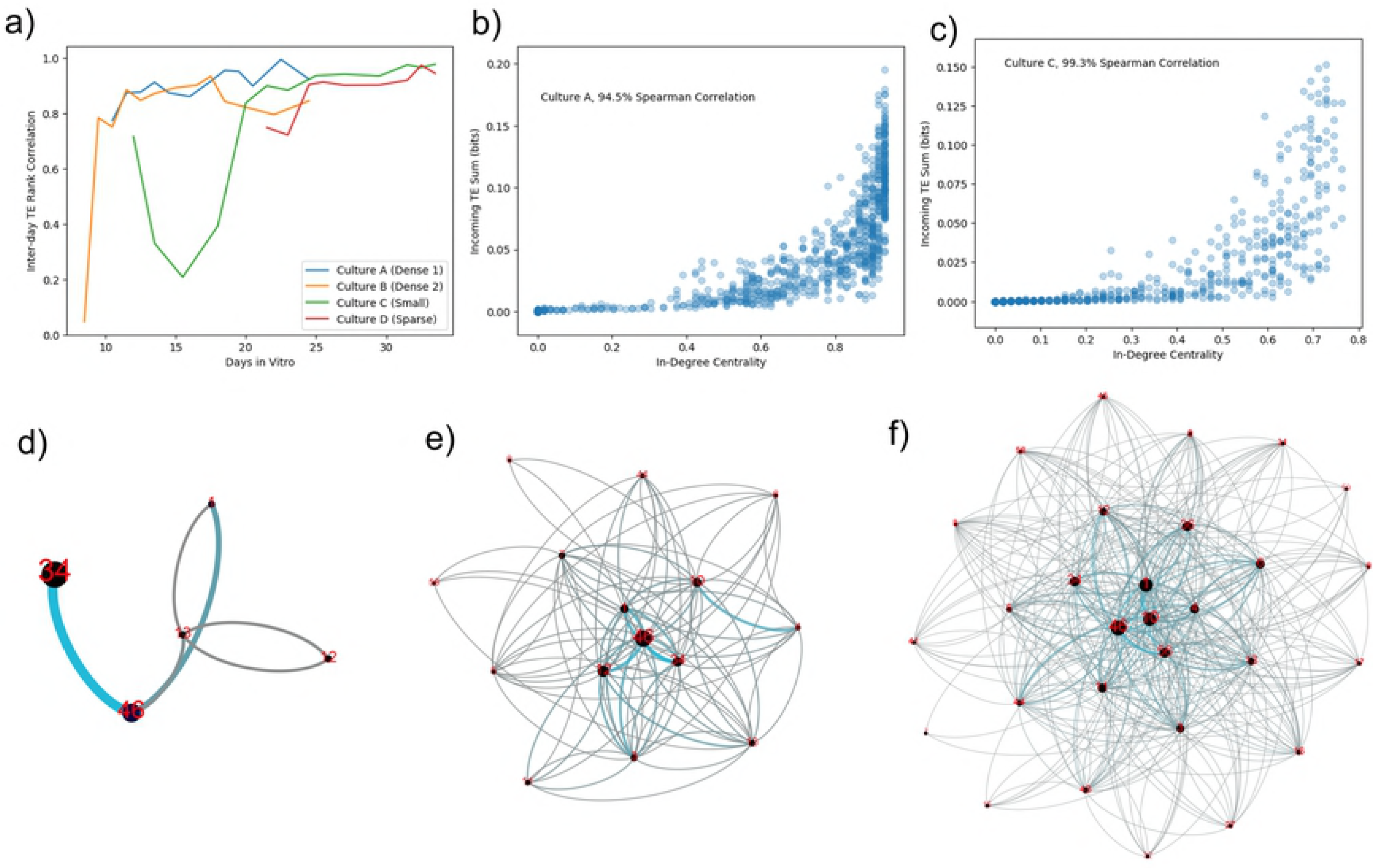
Stability of electrode ranking in R-TE results. a) The correlation of rankings of electrodes by their total incoming TE. A high, consistent correlation suggests this ranking is stable. b,c) The rank correlation of total incoming TE to graphical in-degree, for cultures A and C (respectively). d,e,f) Representations of the connectivity graphs inferred by applying R-TE to recordings of culture D at 21 (d), 25 (e), and 31 (f) DIV. Edge color is proportional to TE value, and node size is proportional to the electrode’s rank of total incoming TE. Nodes which establish connections early (e.g. 1, 34, 46) tend to remain important in the network.

### 2.4 Software Features and Performance

By taking advantage of the sparsity of data cross time and space in MEA recordings, calculation of TE can be greatly sped up. Additionally, surrogate data generation and resampling are independent, and can thus be trivially parallelized. We take advantage of these features to greatly speed up the computation of discrete TE, distributing work across all available resources. As a result, computation times are quite tractable on a modern multi-core machine. Fully analyzing a 60-channel recording, including creating surrogates and resampling values, rarely took more than an hour (Fig 7a).

**Fig.7.**
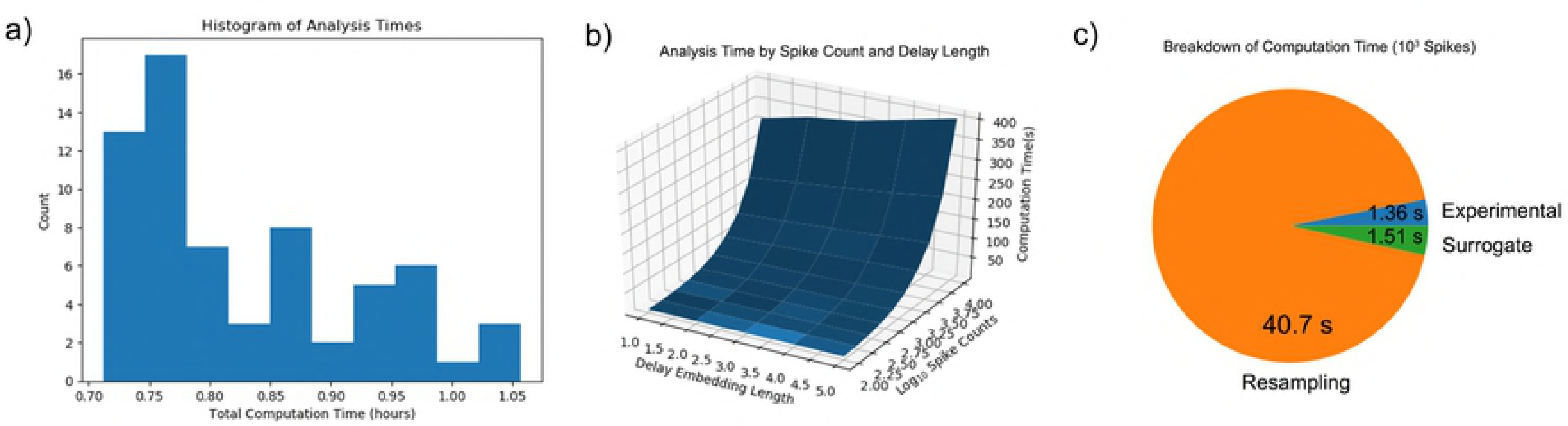
Performance of R-TE. a) An overview of R-TE’s runtime on the MEA recordings, using the specified parameters (w =1 ms, *τ* = 3, 0-20 offsets, 32 resamples, 32 surrogates). b) Runtime scales linearly with spike count, and roughly linearly with delay. c) R-TE’s execution time is dominated by the MCMC resampling process, which must inspect every possible connection separately.

Runtime scales linearly with the number of spikes in the recording, and the length of the delay embedding used (Fig 7b). It is also linearly dependent on the number of surrogates, though this can be offset by parallelizing their generation. Lastly, runtime depends on the number of channels squared. As a consequence, the procedure which dominates the time used is the resampling process (Fig 7c), which must inspect the variability of every possible connection in the recording.

## 3 Conclusion

By applying Markov-chain Monte Carlo (MCMC) methods to inspect the variability of a joint distribution (JD) used to calculate transfer entropy (TE) values, we have developed a method which can capture samples of experimental and surrogate TE values. Non-parametric statistics directly applied to these samples are used as a basis for significance testing, instead of assuming that these values follow a distribution. This creates a more solid basis for statistical significance testing to determine whether a TE value corresponds to a significant connection.

We find that this makes a marked and significant improvement in inferring the connectivity of a neuronal network from its discrete spike train (as recored by an MEA). TE also provides insights into the organization of information flow within this biological network, providing a view into development and identifying the emergence of hubs.

In future work, exporting the resampling algorithm to a GPU computation platform could dramatically increase the speed of computation, as it is a massively parallel problem. Additionally, other methods of determining the optimal TE parameters would be useful to increase confidence the signal is being properly captured.

In conclusion, we believe this work provides a novel basis to infer the functional connectivity of a large neuronal network via transfer entropy, and yields insights into its operation.

## Acknowledgments

We would like to acknowledge Charles Bergeron for providing copies of the spike trains being analyzed.

